# Exposure to bilingual or monolingual maternal speech during pregnancy affects the neurophysiological encoding of speech sounds in neonates differently

**DOI:** 10.1101/2024.02.21.579933

**Authors:** Sonia Arenillas-Alcón, Natàlia Gorina-Careta, Marta Puertollano, Alejandro Mondéjar-Segovia, Siham Ijjou-Kadiri, Jordi Costa-Faidella, María Dolores Gómez-Roig, Carles Escera

**Affiliations:** Brainlab – Cognitive Neuroscience Research Group. Departament de Psicologia Clinica i Psicobiologia, Universitat de Barcelona (Barcelona, Spain); Institut de Neurociènces, Universitat de Barcelona (Barcelona, Spain); Institut de Recerca Sant Joan de Déu, Santa Rosa 39-57, 08950 Esplugues de Llobregat (Catalonia, Spain); BCNatal – Barcelona Center for Maternal Fetal and Neonatal Medicine (Hospital Sant Joan de Déu and Hospital Clínic), University of Barcelona (Catalonia, Spain)

**Keywords:** speech brainstem responses, bilingualism, newborns, early language acquisition

## Abstract

Exposure to maternal speech during the prenatal period shapes speech perception and linguistic preferences, allowing neonates to recognize stories heard frequently in utero and demonstrating an enhanced preference for their mother’s voice and native language. Yet, with a high prevalence of bilingualism worldwide, it remains an open question whether monolingual or bilingual maternal speech during pregnancy influence differently the fetus’ neural mechanisms underlying speech sound encoding. In the present study, the frequency-following response (FFR), an auditory evoked potential that reflects the complex spectrotemporal dynamics of speech sounds, was recorded to a two-vowel /oa/ stimulus in a sample of 131 healthy term neonates within the 1-3 days after birth. Newborns were divided into two groups according to maternal language usage during the last trimester of gestation (monolingual; bilingual). Spectral amplitudes and spectral signal-to-noise ratios (SNR) at the stimulus fundamental (F_0_) and first formant (F_1_) frequencies of each vowel were respectively taken as measures of pitch and formant structure neural encoding. Our results reveal that while spectral amplitudes at F_0_ did not differ between groups, neonates from bilingual mothers exhibited a lower spectral SNR. Additionally, monolingually exposed neonates exhibited a higher spectral amplitude and SNR at F_1_ frequencies. We interpret our results under the consideration that bilingual maternal speech, as compared to monolingual, is characterized by a greater complexity in the speech sound signal, rendering newborns from bilingual mothers more sensitive to a wider range of speech frequencies without generating a particularly strong response at any of them. Our results contribute to an expanding body of research indicating the influence of prenatal experiences on language acquisition and underscore the necessity of including prenatal language exposure in developmental studies on language acquisition, a variable often overlooked yet capable of influencing research outcomes.

## INTRODUCTION

The process of language acquisition has long been a point of uncertainty in research exploring the roots of human language, in particular regarding the open debate of nature versus nurture framework. The nature perspective postulates that language, like other aspects of human knowledge, is primarily determined by genetic factors (Chomsky, 1959; Fodor, 1985; Pinker, 1984), while the nurture viewpoint emphasizes the role of experience in language learning (Piaget, 1977; Tomasello, 2000). Researchers have conducted extensive investigations to understand the initial state and process of language acquisition, providing insights into how environmental and genetic factors interact to fashion language and cognitive function, and the mechanisms underlying brain plasticity (Barkat et al., 2011; Weaver et al., 2004; Werker & Hensch, 2015; Werker & Tees, 2005). It is now widely accepted that both genetic and experiential factors contribute to language acquisition (Gervain & Mehler, 2010; Werker & Curtin, 2005), and researchers are interested in understanding how these factors interact during human development.

Infants at birth already exhibit advanced speech perception and language learning abilities. Newborns manifest a preference for speech over non-speech sounds (Vouloumanos & Werker, 2007), can discriminate between different languages based on their speech rhythms (Ramus et al., 2000), detect word boundaries (Christophe et al., 2001), discriminate words with different patterns of stress (Sansavini et al., 1997), or even distinguish consonant sounds (Cabrera & Gervain, 2020) and encode voice pitch in an adult-like manner (Arenillas-Alcón et al., 2021). These findings support the role of a genetically driven cerebral organization towards processing specific speech characteristics.

However, the prenatal period is not devoid of language experience and the study of its influence on the newborn’s speech and language encoding capacities is receiving increasing attention. Hearing becomes functional and undergoes most of its development around the 26^th^ to 28^th^ week of gestation, allowing the fetus to perceive the maternal speech signal (Anbuhl et al., 2016; Granier-Deferre et al., 2011; May et al., 2011; Moore & Linthicum, 2007; Ruben, 1995). Although the exact characteristics of the acoustic signal reaching the fetus are not fully understood, intrauterine recordings from animal models and simulations suggest that the maternal womb acts as a low-pass filter, attenuating around 30 dB for frequencies over 600-1000 Hz (Gerhardt & Abrams, 2000). The low-frequency components of speech that are transmitted through the uterus include pitch, slow aspects of rhythm and some phonetic information (May et al., 2011). Evidence indicates that prenatal exposure to speech, despite attenuated by the filtering properties of the womb, shapes speech perception and linguistic preferences of newborns, as shown by studies revealing that neonates can recognize a story heard frequently in utero (DeCasper & Spence, 1986), prefer the voice of their mother (DeCasper & Fifer, 1980) and prefer their native language (Moon et al., 1993). Additionally, prenatal learning extends beyond these common preferences. Recent findings indicate that infants acquire specific knowledge of the prosody (Gervain, 2018) and prefer the rhythmic patterns of the language they were exposed to while in utero (Mariani et al., 2023), indicating a very early specialization for their native language.

Yet, with reported rates of bilingualism of around 65% in Europe (Luk, 2017), an open question remains on the influence of prenatal exposure to more than one language on neural plasticity. Over the past 20 years, mounting evidence has suggested that both exposure to a bilingual acoustic environment and learning several languages affect not only language acquisition but a wide range of developmental processes including perception, cognition and brain development (Byers-Heinlein et al., 2019). As bilingual mothers speak using two different sets of phonemic categories and even use two slightly different voice pitch ranges (e.g., Ordin & Mennen, 2017), in-utero bilingual environments are characterized by a greater complexity of the reaching speech signal than monolingual ones. Interestingly, neonates exposed prenatally to a bilingual environment can discriminate their two native languages already at birth and exhibit equal preferences for both (Byers-Heinlein et al., 2010). Thus, it appears clear that linguistic experiences while in utero play a significant role in shaping the early development of speech processing. However, how different prenatal maternal linguistic exposure influences the neural mechanisms underlying speech processing at birth is currently unknown.

A large body of evidence has supported the study of the neural encoding of speech through electrophysiological recordings elicited to speech sounds. In particular, the frequency-following response (FFR) can provide insights into the underlying neural mechanisms associated with prenatal language experience, shedding light on how early linguistic exposure shapes the speech-encoding capacities of newborns. The FFR is an auditory evoked potential elicited by periodic complex sounds that reflects neural synchronization with the auditory eliciting signal along the ascending auditory pathway (Krizman & Kraus, 2019; Skoe & Kraus, 2010), providing an accurate snapshot of the neural encoding of speech sounds. FFR recordings have thus become a useful tool to investigate the ability to distinguish between the pitch of different speakers’ voices and the ability to encode the fine spectrotemporal details that distinguish between different phonemes of speech sounds (Gorina-Careta et al., 2022). The interest in the neonatal FFR arises from its potential to serve as a predictive measure for future language development (Schochat et al., 2017), since alterations in FFR patterns in children have been associated with difficulties in reading and learning, dyslexia, impairments in phonological awareness and even autism (Banai et al., 2009; Basu et al., 2010; Chandrasekaran et al., 2009; Font-Alaminos et al., 2020; Hornickel et al., 2012; King et al., 2002; Lam et al., 2017; Otto-Meyer et al., 2018; Rosenthal, 2020). Interestingly, the FFR reflects the impact of a wide range of auditory experiences in children and adults, including training interventions, musical practice and bilingualism (Carcagno & Plack, 2017; Gorina-Careta et al., 2019; Kraus & Chandrasekaran, 2010; Krizman et al., 2012, 2015; Russo et al., 2005; Skoe et al., 2017; Song et al., 2008). In neonates, FFR recordings have also been used to study the effects at birth of prenatal fetal auditory experiences such as music exposure (Arenillas-Alcón et al., 2023).

In the present study, we aimed to examine the influence of maternal bilingual linguistic exposure in-utero in speech sound encoding at birth. To that end, we recorded FFRs from newborns who had been exposed to either a monolingual or a bilingual fetal environment during the last trimester of gestation and analyzed their capacity to encode voice pitch and vocalic formant structure information.

## METHODS

### Participants

A sample of 131 newborns was recruited from *SJD Barcelona Children’s Hospital* in Barcelona (Spain), and divided into two groups according to the response of their mothers to the question “Did you communicate using more than one language during the last 3 months of pregnancy?” in a language usage questionnaire. Based on the collected responses, a total of 53 newborns were assigned to the group exposed to a monolingual fetal acoustic environment (MON; 27 females; mean gestational age = 39.93 ± 1.03 weeks; mean birth weight = 3321 ± 272 g) and 78 to the bilingual-exposed group (BIL; 34 females; mean gestational age = 39.71 ± 0.99 weeks; mean birth weight = 3327 ± 323 g). No significant differences were found across groups in gestational age (*U* = 1868.500, *p* = 0.351), birth weight (*t* = −0.104, *p* = 0.917) and sex (χ^2^ = 0.686, *p* = 0.408). Importantly, groups did not differ in maternal educational level (χ^2^ = 2.368, *p* = 0.668), a key confounding factor associated with language acquisition and development (Hoff, 2003; Rowe, 2008) closely tied to the linguistic environment a fetus is exposed to. Additionally, we ascertained that groups did not differ in prenatal musical exposure (χ^2^ = 2.019, *p* = 0.364; see Arenillas-Alcón et al. (2023) for details), as it exerts a significant impact on speech encoding capacities at birth (Arenillas-Alcón et al., 2023; Partanen et al., 2022; Partanen, Kujala, Tervaniemi, et al., 2013).

All neonates obtained Apgar scores higher than 8 at 1 and 5 min of life and passed adequately the universal newborn hearing screening (UNHS) before the recruitment. According to the recommendations of the Joint Committee of Infant Hearing (Joint Committee on Infant Hearing, 2019), newborns born from high-risk gestations, after obstetric pathologies or any other kind of risk factors related to hearing impairment were excluded from the recruitment.

Additionally, as performed in previous research from our laboratory (Arenillas-Alcón et al., 2021; Arenillas-Alcón et al., 2023; Ribas-Prats et al., 2019, 2023; Ribas-Prats et al., 2021), both groups of newborns received a standard click-evoked auditory brainstem response (ABR) test to ensure the integrity of the auditory pathway. A click-stimulus, with a duration of 100 µs, was employed during the test, presented at a rate of 19.30 Hz with an intensity of 60 dB sound pressure level (SPL) until a total of 4000 artifact-free repetitions were collected. A prerequisite for participation in the experiment for all newborns was the successful identification of the wave V peak. This study was approved by the Ethical Committee of Clinical Research of the Sant Joan de Déu Foundation (Approval ID: PIC-53-17), and required the mothers to fill out a sociodemographic questionnaire and to sign an informed consent prior to the participation, in line with the Code of Ethics of the World Medical Association (Declaration of Helsinki).

### Stimulus

Neonatal FFRs were collected to a two-vowel stimulus with a rising pitch ending (/oa/; Arenillas-Alcón et al., 2021). The /oa/ stimulus was created in *Praat* (Boersma & Weenink, 2020) and had a total length of 250 ms divided into three different sections, according its fundamental frequency (F_0_) and its formant content (/o/ vowel section: 0-80 ms, F_0_ = 113 Hz, F_1_ = 452 Hz, F_2_ = 791 Hz; /oa/ formant transition section = 80-90 ms; /a/ vowel steady section = 90-160 ms, F_0_ = 113 Hz, F_1_ = 678 Hz, F_2_ = 1017 Hz; /a/ vowel rising section = 160-250 ms, F_0_ = 113-154 Hz, F_1_ = 678 Hz, F_2_ = 1017 Hz; Figure 1A). The /oa/ stimulus was presented at a rate of 3.39 Hz in alternating polarities, and delivered monaurally to the right ear at 60 dB SPL of intensity with an earphone connected to a Flexicoupler disposable adaptor (Natus Medical Incorporated, San Carlos, CA).

**Figure 1. A:**
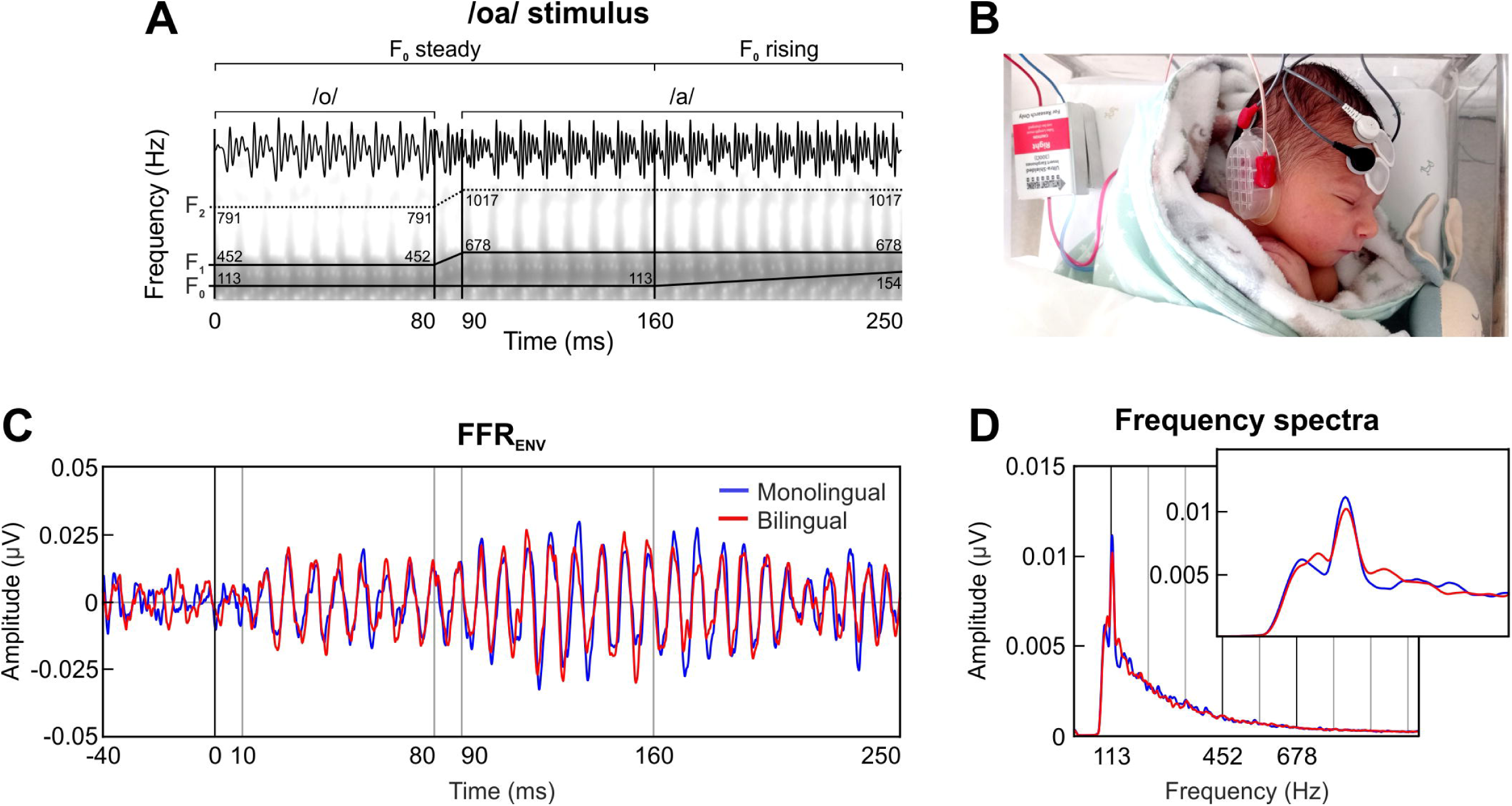
Temporal and spectral representation of the two-vowel auditory stimulus /oa/, with a traces indicating its fundamental frequency (F_0_) and formant structure (F_1_, F_2_). **B**: Recording setup of the three disposable electrodes placed in a vertical montage (active located at Fpz, ground at forehead, references at the right mastoid). Baby’s photo reproduced with the written consent of the neonate’s parents. **C**: Grand-averaged waveform of the FFR_ENV_ in the time domain, retrieved separately for the group exposed to monolingual (blue) and bilingual (red) fetal acoustic environment. **D**: Frequency spectra of the FFR_ENV_ extracted from the steady pitch section of the stimulus (10-160 ms). The inset zooms in a narrower frequency band to illustrate the effect around the F_0_ peak.

### Procedure and data acquisition

After the successful completion of the UNHS, neonates were tested at the hospital room while they were sleeping in their bassinet. Three disposable Ag/AgCl electrodes were placed in a vertical montage configuration (active at Fpz, ground at forehead, reference at the right mastoid, ipsilateral to the auditory stimulation; as shown in Figure 1B), ensuring impedances below 7 kΩ. The presentation of click and speech stimuli was done by using a *SmartEP* platform connected to a *Duet* amplifier, which incorporated the *cABR* and the *Advanced Hearing Research* modules (Intelligent Hearing Systems, Miami, Fl, USA).

The experimental procedure involved the recording of two blocks of click stimuli, followed by four blocks of 1000 artifact-free responses to the /oa/ stimulus. Any electrical activity surpassing ±30 µV threshold was automatically rejected until a total of 4000 presentations was collected. The total mean duration of the recording session was approximately 25 min [2 click blocks x 2000 repetitions x 51.81 ms SOA + 4 /oa/ blocks x 1000 repetitions x 295 ms of stimulus-onset asynchrony (SOA)] including the duration of rejected sweeps. The continuous EEG signal was acquired at a sampling rate of 13333 Hz with an online bandpass filter with cutoff frequencies from 30 to 1500Hz and online epoched from −40.95 ms (pre-stimulus period) to 249.975 ms.

### Data processing and analysis

Data epochs were bandpass filtered offline from 80 to 1500 Hz and averaged separately per stimulus polarity. To highlight the encoding of the stimulus fundamental frequency (F_0_) and to reduce the contribution of cochlear microphonics, neural responses to the two opposite stimulus polarities were added [(Condensation + Rarefaction)/2], obtaining the envelope-following response (FFR_ENV_). Further, to emphasize the FFR components associated with the encoding of the stimulus temporal fine structure, such as the first formant (F_1_), while reducing the impact of envelope-related activity, the neural responses to alternating polarities were subtracted [(Condensation-Rarefaction)/2], yielding the temporal fine structure-following response (FFR_TFS_; Aiken & Picton, 2008; Krizman & Kraus, 2019). Considering the stimulus formant content, we focused our analyses exclusively on the spectral peaks that corresponded to F_1_ frequencies, as F_2_ frequencies fall at the limits of the spectral resolution of the FFR, resulting in elicited neural responses relatively weak and challenging to be accurately observed in newborns (Gorina-Careta et al., 2022). Detailed information regarding the analyzed parameters from the neonatal FFR can be found below. All parameters were computed using custom scripts in Matlab R2019b (The Mathworks, 2019), developed in our laboratory and previously employed in similar analyses in former studies (Arenillas-Alcón et al., 2021).

#### Neural lag

Neural lag served as an indicator of the neural transmission delay within the auditory system, and was assessed to estimate the time passed from cochlear stimulus reception to the onset of neural phase-locking (Arenillas-Alcón et al., 2021; Arenillas-Alcón et al., 2023; Jeng et al., 2010; Liu et al., 2015; Ribas-Prats et al., 2019, 2023; Ribas-Prats et al., 2021). To calculate the neural lag, a cross-correlation analysis was computed between the auditory stimulus and the neural response. The neural lag was determined by identifying the time lag corresponding to the highest cross-correlation value within a time window of 3-13 ms.

#### Pre-stimulus root mean square (RMS) amplitude

The RMS of the pre-stimulus period was employed as a measure of the general magnitude of neural activity over time, and to dismiss electrophysiological disparities in the pre-stimulus region (Arenillas-Alcón et al., 2023; Liu et al., 2015; Ribas-Prats et al., 2019, 2023; Ribas-Prats et al., 2021; White-Schwoch et al., 2015). This measure was computed by squaring each data point within the pre-stimulus region of the neural response (from −40 to 0 ms), calculating the mean of the squared values and subsequently obtaining the square root of the resulting average.

### Voice pitch encoding from FFR_ENV_

#### Spectral amplitude at F_0_

Spectral amplitude at F_0_ (113 Hz) was used as a quantitative measure of the neural phase-locking strength at the specific frequency of interest (Arenillas-Alcón et al., 2021; Arenillas-Alcón et al., 2023; Ribas-Prats et al., 2019, 2023; Ribas-Prats et al., 2021; White-Schwoch et al., 2015). It was computed by applying a fast Fourier transform (FFT; Cooley & Tukey, 1965) to obtain the frequency spectrum of the neural response during the steady pitch section of the stimulus (10-160 ms), and then calculating the average amplitude within a ±5 Hz window centered around the peak of the stimulus F_0_.

#### Signal-to-noise ratio at F_0_

Signal-to-noise ratio (SNR) at F_0_ was analyzed to obtain an estimation of the relative spectral magnitude of the response, taking into account not only to the amplitude value at the F_0_ frequency peak (113 Hz) but also the noise levels at the surrounding frequencies. Therefore, the SNR was calculated by dividing the mean spectral amplitude within a ±5 Hz frequency window centered at the peak of the frequency of interest (113Hz) by the averaged mean amplitude within two additional 28 Hz wide frequency windows (flanks), centered at ±19 Hz from the frequency of interest (80-108 Hz and 118-146 Hz).

### Formant structure encoding from FFR_TFS_

#### Spectral amplitudes at F_1_ peaks

To assess spectral amplitudes at the specific spectral peaks regarding the stimulus F_1_ frequencies (452 Hz [/o/] and 678 Hz [/a/]), the neural responses corresponding to the /o/ section (10–80 ms time window) and the /a/ steady section (90-160 ms time window) were individually analyzed and the respective amplitudes within a ±5 Hz window centered at the peak frequencies corresponding to the vowel formant centers were extracted. The transition from /o/ vowel to /a/ vowel was not analyzed due to its short duration (10 ms).

#### Signal-to-noise ratio at F_1_

To compute the relative spectral magnitude of the response at the stimulus F_1_ frequencies considering noise levels, SNRs at spectral peaks that correspond to the stimulus F_1_ frequencies (452 Hz and 678 Hz) were calculated separately on the /o/ and the /a/-steady sections. To do so, the SNR was calculated by dividing the mean spectral amplitude within a ±5 Hz frequency window centered at the peak of the frequency of interest (452 or 678 Hz) by the averaged mean amplitude within two additional 28 Hz wide frequency windows (flanks), centered at ±26 Hz from the frequency of interest (for 452 Hz peak: 402-430 Hz and 474-502 Hz; for 678 Hz peak: 628-656 Hz and 700-728 Hz).

### Statistical analysis

Statistical analyses were conducted using Jamovi 2.3.26 (The Jamovi Project, 2023). Descriptive statistics were calculated, including the mean, standard deviation (SD), median, first quartile (Q_1_), third quartile (Q_3_), interquartile range (IQR), and minimum and maximum values, for each computed parameter within the two groups of newborns (MON; BIL).

To analyze the effects of prenatal bilingual exposure on neural transmission delay, pre-stimulus root mean square amplitude and voice pitch encoding depending on the normality of the data, two-tailed independent samples *t*-tests or Mann-Whitney U tests were conducted to evaluate significant differences between groups, with Cohen’s *d* being reported as the effect size. Kolmogorov-Smirnov test was used to assess the normal distribution of the samples.

The effects of prenatal bilingual exposure on formant structure encoding were analyzed with two repeated–measures ANOVAs with the factor Stimulus Section (/o/ section; /a/ section) as within-subjects factor and the factor Group (Monolingual; Bilingual) as between-subjects factor for each of the two formant amplitudes (452 and 678 Hz) separately. The Greenhouse–Geisser correction was applied when the assumption of sphericity was violated. Additional two-tailed independent samples Mann-Whitney U post-hoc tests were performed to examine the direction of the effects. Results were considered statistically significant when *p* < 0.05.

## RESULTS

Frequency following responses (FFR) elicited by a two-vowel speech stimulus /oa/ (Figure 1A) were collected from a total sample of 131 newborns divided into two groups according to their prenatal fetal exposure to monolingual (MON) or bilingual (BIL) maternal speech. To comprehensively evaluate the neonates’ ability to encode the pitch and vowel formant structure of speech sounds, the neural responses to the fundamental frequency (F_0_) and the vowels’ first formant (F_1_) were analyzed considering the distinct sound characteristics of the different stimulus sections. All detailed descriptive statistics from the parameters analyzed can be found in Supplementary Table 1.

*Neural transmission delay*. No significant differences were found across groups in neural lag (t_(129)_ = −0.435, *p* = 0.664, Cohen’s *d* = −0.078).

*Pre-stimulus root mean square (RMS) amplitude*. There were no statistically significant differences observed between the groups with regards to the background neural activity preceding the auditory stimulation (U_(129)_ = 1950.000, *p* = 0.585, Rank-biserial correlation = 0.057).

### Voice pitch encoding (FFR_ENV_)

The grand-averaged FFR_ENV_ waveform for each group is illustrated in Figure 1C. To assess the robustness of the voice pitch representation, we analyzed the steady section (10-160 ms) of the /oa/ stimulus with a steady fundamental frequency (F_0_) of 113 Hz.

The grand-averaged spectral representation of the neonatal FFR extracted from each group is depicted in Figure 1D. No differences were found across groups in spectral amplitude at F_0_ computed using the steady pitch section of the stimulus (U_(129)_ = 1809.000, *p* = 0.227, Rank-biserial correlation = 0.125).

Yet, the statistical analyses performed on the F_0_ SNR, which represents the F_0_ relative spectral amplitude in relation with the spectral amplitude of the neighboring frequencies, revealed significant differences between groups, indicating that newborns exposed to a monolingual prenatal fetal environment exhibited significantly larger SNR values as compared to the bilingual exposed neonates (U_(129)_ = 1569.000, *p* = 0.020, Rank-biserial correlation = 0.241).

### Formant structure encoding (FFR_TFS_)

The grand-averaged FFR_TFS_ waveform for each group is shown in Figure 2A. To evaluate the newborns’ ability to encode the formant structure of speech sounds, the /oa/ stimulus included two sections with the same voice pitch but different fine-structure.

**Figure 2.**
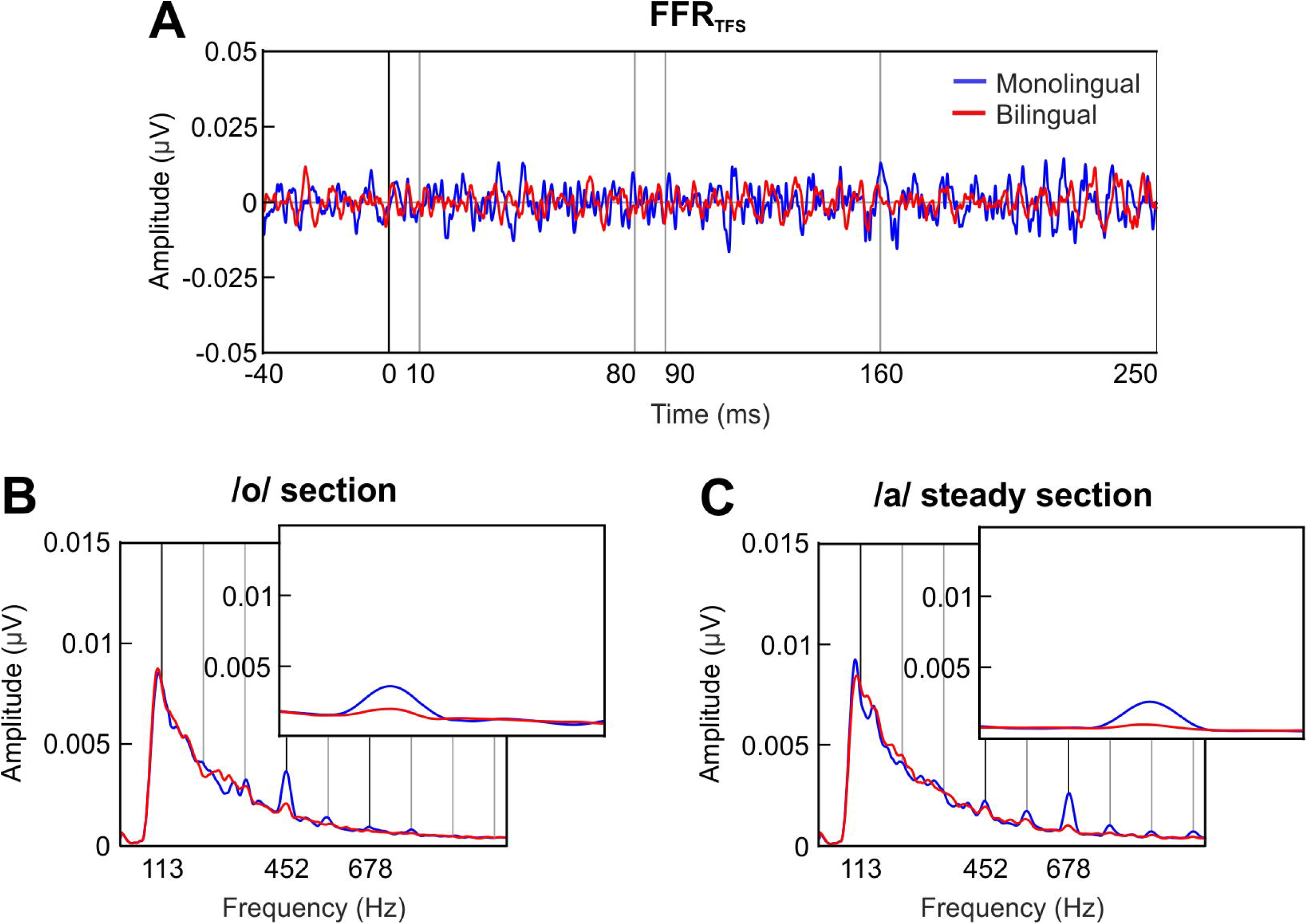
Formant structure encoding. **A:** Grand-averaged waveform of the FFR_TFS_ in the time domain, retrieved separately for the group exposed to a monolingual fetal acoustic environment (blue) and the bilingual-exposed group (red). **B**: Frequency spectra of the FFR_TFS_ extracted from the /o/ section of the stimulus (10-80 ms). The inset zooms in a narrower frequency band to illustrate the effect around the /o/ F_1_ peak (452 Hz) during the /o/ section. **C**: Frequency spectra of the FFR_TFS_ extracted from the /a/ steady section of the stimulus (90-160 ms). The inset zooms in a narrower frequency band to illustrate the effect around the /a/ F_1_ peak (678 Hz) during the /a/ steady section.

Specifically, the /o/ section (10–80 ms) was characterized by a center formant frequency (F_1_) of 452 Hz, and the /a/ steady section (90–160 ms) by a F_1_ frequency of 678 Hz.

Spectral amplitudes were retrieved from the FFR_TFS_ separately from neural responses during the /o/ section and the /a/ steady-pitch section, selecting the spectral peaks corresponding to stimulus F_1_ frequencies.

The grand-averages of the FFR_TFS_ spectral amplitudes during the /o/ section are illustrated in Figure 2B for each group separately, while the spectral representations during the /a/ steady section are depicted in Figure 2C. F_1_ spectral amplitudes during the /o/ section and the /a/ steady section are depicted in Figure 3 for each group at each formant center frequency (452 Hz; 678 Hz) separately.

**Figure 3.**
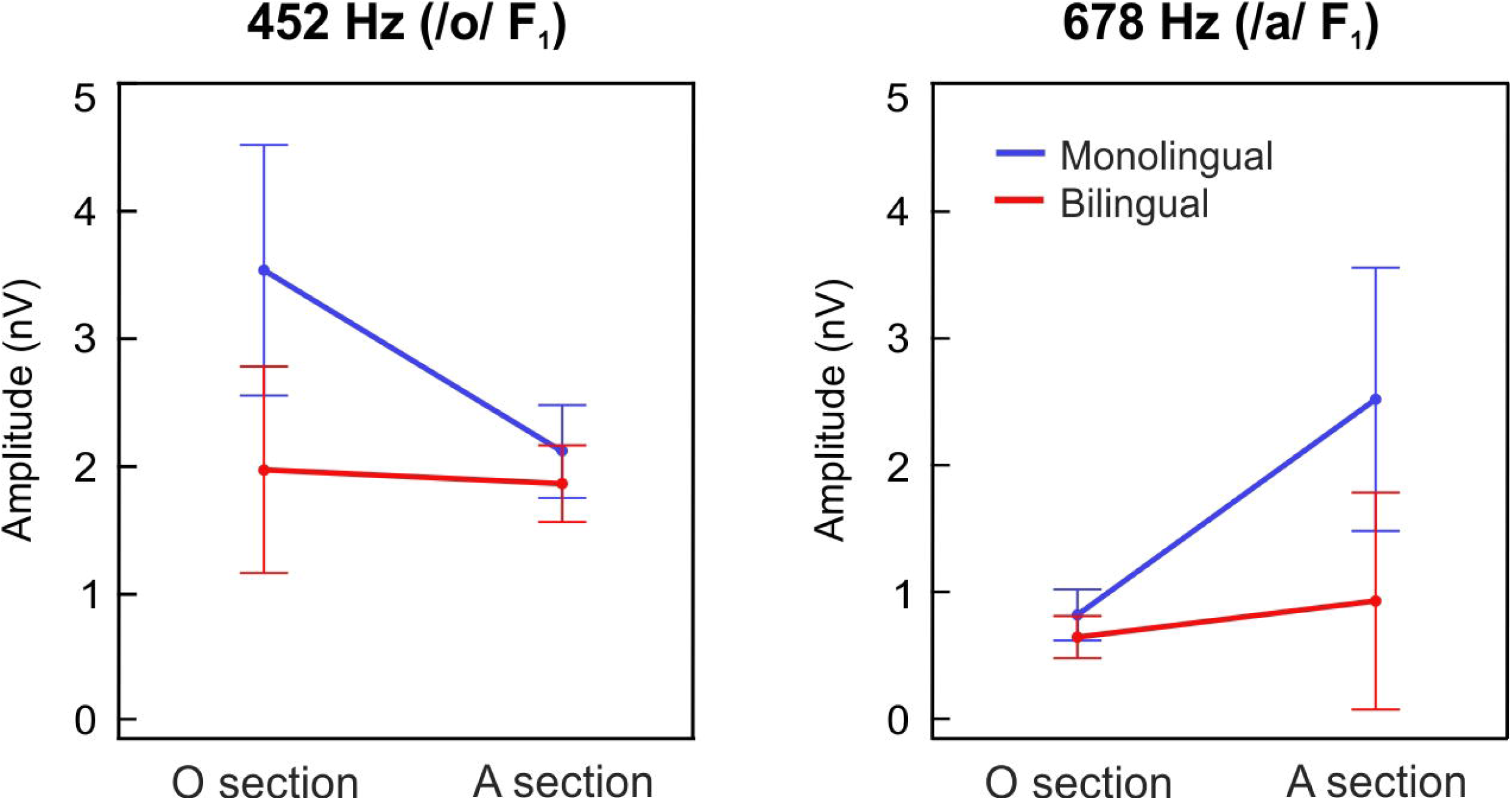
Spectral amplitudes at the first formant (F_1_). F_1_ spectral amplitudes at 452 Hz (left) and 678 Hz (right) during the /o/ section (10-80 ms) and the /a/ steady section (90-160 ms), plotted in blue and red lines for the monolingual and the bilingual-exposed newborns, respectively. Error bars represent 95% confidence intervals.

When analyzing the effects of a prenatal maternal bilingual language exposure in formant spectral amplitude at 452 Hz (Figure 3, left panel), which corresponds to the F_1_ center frequency of the /o/ vowel, a main effect of group revealed significantly greater spectral amplitudes in the MON group as compared to the BIL (group main effect; F_(1,129)_ = 5.115, *p* = 0.025, ηp2 = 0.038). Moreover, a significantly larger spectral amplitude was observed during the /o/ section vs. /a/ steady section (stimulus section main effect; F_(1,129)_ = 7.910, *p* = 0.006, ηp2 = 0.058), thus indicating a proper encoding of the vowel /o/ in its corresponding stimulus section. Interestingly, a significant interaction of group per stimulus section was identified as well (interaction; F_(1,129)_ = 5.817, *p* = 0.017, ηp2 = 0.043), demonstrating that MON neonates showed significantly larger spectral amplitudes during the /o/ section at its corresponding formant frequency than BIL.

Similar results were observed when analyzing the effects of a prenatal maternal bilingual language exposure in the formant encoding at 678Hz (Figure 3, right panel), which corresponds to the F_1_ center frequency of the /a/ vowel. A main effect of group revealed significantly greater spectral amplitudes in the MON group as compared to the BIL (group main effect; F_(1,129)_ = 5.077, *p* = 0.026, ηp2 = 0.038). Moreover, a significantly larger spectral amplitude at 678 Hz during the /a/ steady section vs. /o/ section was observed (stimulus section main effect; F_(1,129)_ = 11.412, *p* < 0.001, ηp2 = 0.081), thus indicating a proper encoding of the /a/ vowel in its corresponding stimulus section. Interestingly, a significant interaction of group per stimulus section was also identified (interaction; F_(1,129)_ = 5.812, *p* = 0.017, ηp2 = 0.043), demonstrating that the MON group exhibited higher spectral amplitudes during the /a/ steady section at its corresponding frequency than the BIL.

The same pattern of results was obtained when comparing the relative spectral amplitude of the response at the stimulus F_1_ frequencies taking into account the neural response to the neighboring frequencies. When analyzing the effects of a fetal maternal bilingual language exposure in SNR at 452 Hz, which corresponds to the F_1_ of the /o/ vowel, a main effect of group revealed significantly greater spectral amplitudes in the MON group as compared to the BIL (group main effect; F_(1,129)_ = 8.300, *p* = 0.005, ηp2 = 0.060). Moreover, a significantly larger spectral amplitude was observed during the o/ section vs. /a/ steady section (stimulus section main effect; F_(1,129)_ = 7.910, *p* = 0.006, ηp2 = 0.058). A significant interaction of group per stimulus section was identified as well (interaction; F_(1,129)_ = 7.30, *p* = 0.008, ηp2 = 0.054).

Similar effects were observed when analyzing the effects of a prenatal bilingual environment in the formant SNR at 678 Hz, which corresponds to the frequency of the /a/ vowel. A main effect of group revealed significantly greater spectral amplitudes in the MON group as compared to the BIL (group main effect; F_(1,129)_ = 7.01, *p* = 0.009, ηp2 = 0.052). Moreover, a significantly larger spectral amplitude at 678 Hz during the /a/ steady section vs. /o/ section was observed (stimulus section main effect; F_(1,129)_ = 23.300, *p* < 0.001, ηp2 = 0.153). Finally, a significant interaction of group per stimulus section was also identified (interaction; F_(1,129)_ = 10.000, *p* = 0.002, ηp2 = 0.072).

## DISCUSSION

The present study investigated the impact of maternal bilingual speech during pregnancy on the neural encoding of speech pitch and vowel formant structure in neonates. A total sample of 131 healthy-term newborns was divided into two groups according to their monolingual or bilingual prenatal exposure during the last trimester of gestation, as reported by their mothers through a questionnaire. FFRs elicited to a two-vowel speech stimulus /oa/ (Arenillas-Alcón et al., 2021) were recorded to assess the neural responses to the stimulus’ fundamental frequency (F_0_ = 113 Hz; related to voice pitch encoding) and the first formant of each vowel (/o/ F_1_ = 452 Hz; /a/ F_1_ = 678 Hz; related to vowel formant structure encoding). Our results revealed that the neural representation of pitch, as indexed by the spectral amplitude of the FFR_ENV_ at the stimulus F_0_, did not differ between monolingual and bilingual exposure groups, but bilingually exposed neonates exhibited a lower signal-to-noise ratio (SNR) at the F_0_ spectral peak, suggesting the contribution of a higher spectral noise at neighboring frequencies. Additionally, monolingually exposed neonates exhibited larger spectral amplitudes and SNRs of the FFR_TFS_ at the formant peak frequencies (F_1_) of the speech stimulus used, indicating a stronger encoding of vocalic structure. Furthermore, no significant group differences were observed in neural lag and pre-stimulus root mean square (RMS) amplitude, implying comparable neural transmission delays and absence of a distinct overall neural activity prior to the auditory stimulation. Together, these findings provide novel insights into the effects of prenatal language exposure on the neural encoding of speech sounds at birth.

Pitch is a crucial attribute in the perception of periodic speech sounds, as it conveys prosodic information, facilitates speaker recognition and speech segmentation, accelerates phoneme acquisition in tonal languages, helps with language comprehension in noisy environments and even contributes to the perception of the emotional state in a conversation (Arenillas-Alcón et al., 2021; Benavides-Varela et al., 2012; Cabrera & Gervain, 2020; Gervain, 2018; Musacchia et al., 2007; Partanen, Kujala, Näätänen, et al., 2013; Plack et al., 2014; Ribas-Prats et al., 2021). The fact that neural mechanisms underlying voice pitch encoding are already mature at birth (Arenillas-Alcón et al., 2021; Cabrera & Gervain, 2020; Jeng et al., 2011; Ribas-Prats et al., 2019) suggests that pitch may play a crucial role in the very first stages of language acquisition (Jeng et al., 2016). Pitch could provide a neural synchrony channel onto which separate neural representations of other speech features would anchor as parts of an ensemble that would, ultimately, give rise to a coherent percept (Eggermont, 2001).

Previous studies demonstrated that pitch and pitch contour discrimination drastically improve with training (e.g., Carcagno & Plack, 2017). In this regard, growing up in a bilingual environment, which is characterized as more demanding, dynamic, phonologically rich and requiring heightened attention to all linguistic input, is related to a strengthened neural representation of pitch (Krizman et al., 2012, 2015). Different languages have distinct overall height pitch levels. For example, Polish was found to have a higher pitch compared to American English (Majewski et al., 1972); Mandarin, a higher pitch than English (Keating & Kuo, 2012); Japanese, a higher pitch than Dutch (Van Bezooijen, 1995); or Slavic languages, a higher pitch than Germanic ones (Andreeva et al., 2014). Further, speakers of two phonologically similar dialects exhibit differences in their height pitch levels (e.g., two different dialects of Mandarin; Deutsch et al., 2009). With continued exposure to these complex linguistic contexts, the auditory system gradually becomes finely tuned to process sound more efficiently (Krizman et al., 2012). Thus, individuals with years of exposure and interaction with bilingual environments develop enhanced flexibility and speech-encoding abilities. Most notably, previous studies have shown that bilingual individuals, particularly females, exhibit different pitch frequency ranges depending on the language they speak (Ordin & Mennen, 2017). As speech F_0_ is an acoustic feature reliably transmitted through the womb (Gerhardt & Abrams, 2000; May et al., 2011), bilingual mothers provide their children with a higher pitch variability in utero.

Considering the reviewed literature, if the developing auditory system of a fetus, who underwent approximately three months of noninteractional exposure to degraded speech, responded to acoustic exposure as the mature one, we would expect newborns from bilingual mothers to exhibit a higher neural encoding of voice pitch. But our results showed otherwise. We found no differences across groups in FFR_ENV_ spectral amplitudes at F_0_, which aligns with the idea that pitch processing mechanisms are already mature at birth. Yet, we observed a decreased SNR at the F_0_ in newborns who were prenatally exposed to a bilingual environment. We attempt to reconcile our seemingly contradicting results by hypothesizing that the higher spectral amplitudes found in bilingually exposed neonates at F_0_ neighbouring frequencies reflect an increased sensitivity to a wider range of pitch frequencies without yet generating a particularly strong response at any of them. This view aligns with research about perceptual phonetic development showing broad phoneme discrimination capabilities in newborns as compared with the more language-specific discrimination abilities described in older infants and adult participants (Aslin et al., 1981; Gervain & Werker, 2008; Saffran et al., 2006), especially when growing in bilingual environments. All in all, our findings could potentially be explained by developmental differences linked to exposure times.

Our results also reveal a modulation of the neural encoding of vowel formants (F_1_) depending on prenatal linguistic exposure. In particular, monolingual-exposed neonates exhibited higher spectral amplitudes at the corresponding formant frequencies of the stimulus’ /o/ and steady-/a/ vowels. In a previous study, we found that while the neural encoding of pitch was adult-like at birth, formant encoding was still immature (Arenillas-Alcón et al., 2021). As vowel formant center frequencies are language specific and stable regardless of voice pitch variation, which also presents slight modulations in monolingual individuals during natural speaking, the auditory system of a monolingual-exposed fetus receives a more consistent and less complex phonetic repertoire than that of a bilingual-exposed. This would possibly lead to a more effective and accurate encoding of the specific language vowel sound characteristics at birth. Simply put, monolingual newborns seem to have an advantage in processing the specific sounds of their mother tongue, a finding previously attributed to postnatal linguistic exposure (Kuhl, 2010). Our findings thus highlight the greater variability of acoustic speech inputs to which the fetus of bilingual mothers would be exposed and therefore suggest the need for bilinguals to develop a different phonological representation for each of the languages (Sebastian-Gallés et al., 2006). Further investigation into the developmental trajectories of auditory processing in different populations of newborns, with different prenatal auditory experiences, and using language-specific phonetic contrasts (e.g., Catalan contrasts such as /e - ɛ/), which are especially difficult –when not impossible– to detect for Spanish-monolinguals (Pallier et al., 1997, 2001), may shed more light on this issue.

Overall, our findings emphasize the potential importance of prenatal linguistic exposure in shaping the neural mechanisms underlying language acquisition and highlight the sensitivity of the FFR in capturing these subtle changes. The results add to a growing body of research that suggests a role for prenatal fetal experiences in modeling language acquisition (Arenillas-Alcón et al., 2023; Gervain, 2015, 2018; Moon et al., 2012; Partanen, Kujala, Tervaniemi, et al., 2013). Furthermore, they also highlight the importance of considering prenatal language exposure in developmental studies about language acquisition, a factor that is not routinely measured and reported, and that may contribute to divergent findings.

## CONCLUSION

Overall, the present study contributes significant insights into the impact of prenatal bilingual exposure on the neural encoding of speech sounds at birth, thereby increasing our knowledge of the early stages of language acquisition. The observed differences in the encoding of voice pitch and formant structure depending on prenatal linguistic exposure highlight the remarkable plasticity and learning potential of the human brain even before birth, emphasizing the complex interaction between genetic and environmental factors in shaping our cognitive abilities and linguistic development.

## Acknowledgements

The authors would like to thank to all families who selflessly participated in this study and made it possible.

## AUTHOR CONTRIBUTIONS

**S.A.A.:** Conceptualization, Methodology, Investigation, Formal analysis, Writing-Original draft, Editing, Visualization. **N.G.C:** Conceptualization, Methodology, Investigation, Formal analysis, Writing-Original draft, Writing-Review & Editing. **M.P.:** Methodology, Investigation. **A.M.S.:** Methodology, Investigation. **S.I.K.:** Methodology, Investigation. **J.C.F.:** Conceptualization, Methodology, Writing-Review & Editing, Supervision. **M.D.G.R.:** Funding acquisition, Resources. **C.E.:** Conceptualization, Methodology, Writing-Review & Editing, Funding acquisition, Resources, Supervision.

## REFERENCES

Aiken, S. J., & Picton, T. W. (2008). Envelope and spectral frequency-following responses to vowel sounds. Hearing Research, 245(1–2), 35–47. 10.1016/j.heares.2008.08.004

Anbuhl, K. L., Uhler, K. M., Werner, L. A., & Tollin, D. J. (2016). Early Development of the Human Auditory System. In R. A. Polin, S. H. Abman, D. Rowitch, & W. E. Benitz (Eds.), Fetal and Neonatal Physiology (5th ed., pp. 1396–1410). Elsevier. 10.1016/B978-0-323-35214-7.00138-4

Andreeva, B., Demenko, G., Möbius, B., Zimmerer, F., Jügler, J., & Jastrzebska, M. (2014). Differences of pitch profiles in Germanic and Slavic languages. In Proceedings of Interspeech. ISCA.

Arenillas-Alcón, S., Costa-Faidella, J., Ribas-Prats, T., Gómez-Roig, M. D., & Escera, C. (2021). Neural encoding of voice pitch and formant structure at birth as revealed by frequency-following responses. Scientific Reports, 11, 6660. 10.1038/s41598-021-85799-x

Arenillas-Alcón, S., Ribas-Prats, T., Puertollano, M., Mondéjar-Segovia, A., Gómez-Roig, M. D., Costa-Faidella, J., & Escera, C. (2023). Prenatal daily musical exposure is associated with enhanced neural representation of speech fundamental frequency: Evidence from neonatal frequency-following responses. *Developmental Science*, e13362. 10.1111/desc.13362

Aslin, R. N., Pisoni, D. B., Hennessy, B. L., & Perey, A. J. (1981). Discrimination of voice onset time by human infants: new findings and implications for the effects of early experience. Child Development, 52, 1135–1145.

Banai, K., Hornickel, J., Skoe, E., Nicol, T., Zecker, S., & Kraus, N. (2009). Reading and Subcortical Auditory Function. Cerebral Cortex, 19, 2699–2707. 10.1093/cercor/bhp024

Barkat, T., Polley, D., & Hensch, T. (2011). A critical period for auditory thalamocortical connectivity. Nature Neuroscience, 14, 1189–1194.

Basu, M., Krishnan, A., & Weber-Fox, C. (2010). Brainstem correlates of temporal auditory processing in children with specific language impairment. Developmental Science, 13(1), 77–91. 10.1111/j.1467-7687.2009.00849.x

Benavides-Varela, S., Hochmann, J. R., Macagno, F., Nespor, M., & Mehler, J. (2012). Newborn’s brain activity signals the origin of word memories. Proceedings of the National Academy of Sciences of the United States of America, 109(44), 17908– 17913. 10.1073/pnas.1205413109

Boersma, P., & Weenink, D. (2020). Praat: doing phonetics by computer (Version 6.1.09; p. Version 6.1.09). http://www.praat.org/

Byers-Heinlein, K., Burns, T. C., & Werker, J. F. (2010). The Roots of Bilingualism in Newborns. Psychological Science, 21(3), 343–348. 10.1177/0956797609360758

Byers-Heinlein, K., Esposito, A. G., Winsler, A., Marian, V., Castro, D. C., & Luk, G. (2019). The Case for Measuring and Reporting Bilingualism in Developmental Research. Collabra Psychology, 5(1), 37. 10.1525/collabra.233.The

Cabrera, L., & Gervain, J. (2020). Speech perception at birth: The brain encodes fast and slow temporal information. Science Advances, 6(30), eaba7830. 10.1126/sciadv.aba7830

Carcagno, S., & Plack, C. J. (2017). Short-Term Learning and Memory: Training and Perceptual Learning. In N. Kraus, S. Anderson, T. White-Schwoch, R. R. Fay, & A. N. Popper (Eds.), The Frequency-Following Response: A Window into Human Communication (Vol. 61, pp. 75–100). Springer Nature.

Chandrasekaran, B., Hornickel, J., Skoe, E., Nicol, T., & Kraus, N. (2009). Context-dependent encoding in the human auditory brainstem relates to hearing speech in noise: Implications for developmental dyslexia. Neuron, 64(3), 311–319. 10.1016/j.neuron.2009.10.006

Chomsky, N. (1959). A review of B.F. Skinner’s verbal behavior. Language, 35, 26–58.

Christophe, A., Mehler, J., & Sebastian-Galles, N. (2001). Perception of prosodic boundary correlates by newborn infants. Infancy, 2, 358–394.

Cooley, J. W., & Tukey, J. W. (1965). An Algorithm for the Machine Calculation of Complex Fourier Series. Mathematics of Computation, 19(90), 297.

Corp., I. (n.d.). SPSS 25.0.

DeCasper, A. J., & Fifer, W. (1980). Of human bonding: newborns prefer their mothers’ voice. Science, 208(4448), 1174–1176. 10.1126/science.7375928

DeCasper, A. J., & Spence, M. J. (1986). Prenatal maternal speech influences newborns’ perception of speech sounds. Infant Behavior & Development, 9(2), 133–150. 10.1016/0163-6383(86)90025-1

Deutsch, D., Le, J., Shen, J., & Henthorn, T. (2009). The pitch levels of female speech in two Chinese villages. The Journal of the Acoustical Society of America, 125, EL208–EL213.

Eggermont, J. J. (2001). Between sound and perception: reviewing the search for a neural code. Hearing Research, 157(1–2), 1–42. 10.1016/S0378-5955(01)00259-3

Fodor, J. A. (1985). Precis of the modularity of mind. Behavioral and Brain Sciences, 8, 1–42.

Font-Alaminos, M., Cornella, M., Costa-Faidella, J., Hervás, A., Leung, S., Rueda, I., & Escera, C. (2020). Increased subcortical neural responses to repeating auditory stimulation in children with autism spectrum disorder. Biological Psychology, 149, 107807. 10.1016/j.biopsycho.2019.107807

Gerhardt, K. J., & Abrams, R. M. (2000). Fetal Exposures to Sound and Vibroacoustic Stimulation. Journal of Perinatology, 20, 20–29.

Gervain, J. (2015). Plasticity in early language acquisition: the effects of prenatal and early childhood experience. Current Opinion in Neurobiology, 35, 13–20. 10.1016/j.conb.2015.05.004

Gervain, J. (2018). The role of prenatal experience in language development. Current Opinion in Behavioral Sciences, 21, 62–67. 10.1016/j.cobeha.2018.02.004

Gervain, J., & Mehler, J. (2010). Speech Perception and Language Acquisition in the First Year of Life. Annual Review of Psychology, 61, 191–218. 10.1146/annurev.psych.093008.100408

Gervain, J., & Werker, J. F. (2008). How Infant Speech Perception Contributes to Language Acquisition. Language and Linguistics Compass, 2(6), 1149–1170. 10.1111/j.1749-818X.2008.00089.x

Gorina-Careta, N., Ribas-Prats, T., Arenillas-Alcón, S., Puertollano, M., Gómez-Roig, M. D., & Escera, C. (2022). Neonatal Frequency-Following Responses: A Methodological Framework for Clinical Applications. Seminars in Hearing, 43(3), 162–176. 10.1055/s-0042-1756162

Gorina-Careta, N., Ribas-Prats, T., Costa-Faidella, J., & Escera, C. (2019). Auditory Frequency-Following Responses. In D. Jaeger & R. Jung (Eds.), Encyclopedia of Computational Neuroscience (pp. 1–13). Springer, New York, NY. 10.1007/978-1-4614-7320-6_100689-1

Granier-Deferre, C., Ribeiro, A., Jacquet, A. Y., & Bassereau, S. (2011). Near-term fetuses process temporal features of speech. Developmental Science, 14(2), 336–352. 10.1111/j.1467-7687.2010.00978.x

Hoff, E. (2003). The Specificity of Environmental Influence: Socioeconomic Status Affects Early Vocabulary Development Via Maternal Speech. Child Development, 74(5), 1368–1378. 10.1111/1467-8624.00612

Hornickel, J., Anderson, S., Skoe, E., Yi, H.-G., & Kraus, N. (2012). Subcortical representation of speech fine structure relates to reading ability. NeuroReport, 23(1), 6–9. 10.1097/WNR.0b013e32834d2ffd

Jeng, F. C., Hu, J., Dickman, B., Montgomery-Reagan, K., Tong, M., Wu, G., & Lin, C. Der. (2011). Cross-linguistic comparison of frequency-following responses to voice pitch in american and chinese neonates and adults. Ear and Hearing, 32(6), 699–707. 10.1097/AUD.0b013e31821cc0df

Jeng, F. C., Lin, C.-D., & Wang, T.-C. (2016). Subcortical neural representation to Mandarin pitch contours in American and Chinese newborns. The Journal of the Acoustical Society of America, 139(6), 190–195. 10.1121/1.4953998

Jeng, F. C., Schnabel, E. A., Dickman, B. M., Hu, J., Li, X., Lin, C.-D., & Chung, H.-K. (2010). Early Maturation of Frequency-Following Responses to Voice Pitch in Infants with Normal Hearing. Perceptual and Motor Skills, 111(3), 765–784. 10.2466/10.22.24.PMS.111.6.765-784

Joint Committee on Infant Hearing. (2019). Year 2019 Position Statement: Principles and Guidelines for Early Hearing Detection and Intervention Programs. Journal of Early Hearing Detection and Intervention, 4(2), 1–44. 10.1542/peds.106.4.798

Keating, P., & Kuo, G. (2012). Comparison of speaking fundamental frequency in English and Mandarin. The Journal of the Acoustical Society of America, 132, 1050–1060.

King, C., Warrier, C. M., Hayes, E., & Kraus, N. (2002). Deficits in auditory brainstem pathway encoding of speech sounds in children with learning problems. Neuroscience Letters, 319, 111–115. 10.1016/S0304-3940(01)02556-3

Kraus, N., & Chandrasekaran, B. (2010). Music training for the development of auditory skills. Nature Reviews Neuroscience, 11, 599–605. 10.1038/nrn2882

Krizman, J., & Kraus, N. (2019). Analyzing the FFR: A tutorial for decoding the richness of auditory function. Hearing Research, 382, 166–174. 10.1016/j.heares.2019.107779

Krizman, J., Marian, V., Shook, A., Skoe, E., & Kraus, N. (2012). Subcortical encoding of sound is enhanced in bilinguals and relates to executive function advantages. PNAS, 109(20), 7877–7881. 10.1073/pnas.1201575109

Krizman, J., Slater, J., Skoe, E., Marian, V., & Kraus, N. (2015). Neural processing of speech in children is influenced by bilingual experience. Neuroscience Letters, 0, 48–53. 10.1016/j.neulet.2014.11.011.Neural

Kuhl, P. K. (2010). Brain Mechanisms in Early Language Acquisition. Neuron, 67(5), 713–727. 10.1016/j.cortex.2009.08.003.Predictive

Lam, S. S.-Y., White-Schwoch, T., Zecker, S. G., Hornickel, J., & Kraus, N. (2017). Neural stability: A reflection of automaticity in reading. Neuropsychologia, 103, 162–167. 10.1016/j.physbeh.2017.03.040

Liu, F., Maggu, A. R., Lau, J. C. Y., & Wong, P. C. M. (2015). Brainstem encoding of speech and musical stimuli in congenital amusia: evidence from Cantonese speakers. Frontiers in Human Neuroscience, 8, 1–19. 10.3389/fnhum.2014.01029

Luk, G. (2017). Bilingualism. In B. Hopkins, E. Geangu, & S. Linkenauger (Eds.), The Cambridge Encyclopedia of Child Development (2nd ed., pp. 385–391). Cambridge University. 10.1017/9781316216491.062

Majewski, W., Hollien, H., & Zalewski, J. (1972). Speaking fundamental frequency of Polish adult males. Phonetica, 25, 119–125.

Mariani, B., Nicoletti, G., Barzon, G., Ortiz-Barajas, M. C., Shukla, M., Guevara, R., Suweis, S. S., & Gervain, J. (2023). Prenatal experience with language shapes the brain. Science Advances, 9(47), eadj3524. 10.1126/sciadv.adj3524

May, L., Byers-Heinlein, K., Gervain, J., & Werker, J. F. (2011). Language and the newborn brain: does prenatal language experience shape the neonate neural response to speech? Frontiers in Psychology, 2(222), 1–9. 10.3389/fpsyg.2011.00222

Moon, C., Cooper, R. P., & Fifer, W. P. (1993). Two-day-olds prefer their native language. Infant Behavior and Development, 16(4), 495–500.

Moon, C., Lagercrantz, H., & Kuhl, P. K. (2012). Language experienced in utero affects vowel perception after birth: a two-country study. Acta Paedriatrica, 102, 156–160. 10.1111/apa.12098

Moore, J. K., & Linthicum, F. H. (2007). The human auditory system: A timeline of development. International Journal of Audiology, 46(9), 460–478. 10.1080/14992020701383019

Musacchia, G., Sams, M., Skoe, E., & Kraus, N. (2007). Musicians have enhanced subcortical auditory and audiovisual processing of speech and music. Proceedings of the National Academy of Sciences of the United States of America, 104(40), 15894–15898. 10.1073/pnas.0701498104

Ordin, M., & Mennen, I. (2017). Cross-linguistic differences in bilinguals’ fundamental frequency ranges. Journal of Speech, Language, and Hearing Research, 60, 1493– 1506. 10.1044/2016_JSLHR-S-16-0315

Otto-Meyer, S., Krizman, J., White-Schwoch, T., & Kraus, N. (2018). Children with autism spectrum disorder have unstable neural responses to sound. Experimental Brain Research, 236, 733–743. 10.1007/s00221-017-5164-4

Pallier, C., Bosch, L., & Sebastián-Gallés, N. (1997). A limit on behavioral plasticity in vowel acquisition. Cognition, 64(3), B9–B17. 10.1016/s0010-0277(97)00030-9

Pallier, C., Colomé, A., & Sebastian-Gallés, N. (2001). The Influence of Native-Language Phonology on Lexical Access: Exemplar-Based versus Abstract Lexical Entries. Psychological Science, 12(6), 445–449. 10.1111/1467-9280.00383

Partanen, E., Kujala, T., Näätänen, R., Liitola, A., Sambeth, A., & Huotilainen, M. (2013). Learning-induced neural plasticity of speech processing before birth. PNAS, 110(37), 15145–15150. 10.1073/pnas.1302159110

Partanen, E., Kujala, T., Tervaniemi, M., & Huotilainen, M. (2013). Prenatal music exposure induces long-term neural effects. PLoS ONE, 8(10), e78946. 10.1371/journal.pone.0078946

Partanen, E., Mårtensson, G., Hugoson, P., Huotilainen, M., Fellman, V., & Ådén, U. (2022). Auditory Processing of the Brain Is Enhanced by Parental Singing for Preterm Infants. Frontiers in Neural Circuits, 16, 772008. 10.3389/fnins.2022.772008

Piaget, J. (1977). The language and thought of the child. In The Essential Piaget (pp. 66–83). Basic Books Inc.

Pinker, S. (1984). Language Learnability and Language Development. Harvard University Press.

Plack, C. J., Barker, D., & Hall, D. A. (2014). Pitch coding and pitch processing in the human brain. Hearing Research, 307, 53–64. 10.1016/j.heares.2013.07.020

Ramus, F., Hauser, M., Miller, C., Morris, D., & Mehler, J. (2000). Language discrimination by human newborns and by cotton-top tamarin monkeys. Science, 288, 349–351.

Ribas-Prats, T., Almeida, L., Costa-Faidella, J., Plana, M., Corral, M. J., Gómez-Roig, M. D., & Escera, C. (2019). The frequency-following response (FFR) to speech stimuli: A normative dataset in healthy newborns. Hearing Research, 371, 28–39. 10.1016/j.heares.2018.11.001

Ribas-Prats, T., Arenillas-Alcón, S., Pérez-Cruz, M., Costa-Faidella, J., Gómez-Roig, M. D., & Escera, C. (2023). Speech-encoding deficits in neonates born large-for-gestational age as revealed with the frequency-following response. Ear and Hearing, 44(4), 829–841. 10.1097/AUD.0000000000001330

Ribas-Prats, T., Arenillas-Alcón, S., Lip-Sosa, D. L., Costa-Faidella, J., Mazarico, E., Gómez-Roig, M. D., & Escera, C. (2021). Deficient neural encoding of speech sounds in term neonates born after fetal growth restriction. Developmental Science, 1–15. 10.1111/desc.13189

Rosenthal, M. A. (2020). A systematic review of the voice-tagging hypothesis of speech-in-noise perception. Neuropsychologia, 136(November 2019), 107256. 10.1016/j.neuropsychologia.2019.107256

Rowe, M. (2008). Child-directed speech: relation to socioeconomic status, knowledge of child development and child vocabulary skill. Journal of Child Language, 35(1), 185–205.

Ruben, R. J. (1995). The ontogeny of human hearing. International Journal of Pediatric Otorhinolaryngology, 32, 199–204. 10.1016/0165-5876(94)01159-U

Russo, N. M., Nicol, T. G., Zecker, S. G., Hayes, E. A., & Kraus, N. (2005). Auditory training improves neural timing in the human brainstem. Behavioural Brain Research, 156, 95–103. 10.1016/j.bbr.2004.05.012

Saffran, J. R., Werker, J. F., & Werner, L. A. (2006). The Infant’s Auditory World: Hearing, Speech, and the Beginnings of Language. In D. Kuhn, R. S. Siegler, W. Damon, & R. M. Lerner (Eds.), Handbook of child psychology: Cognition, perception, and language (pp. 58–108). John Wiley & Sons, Inc.

Sansavini, A., Bertoncini, J., & Giovanelli, G. (1997). Newborns discriminate the rhythm of multisyllabic stressed words. Developmental Psychology, 33, 3–11.

Schochat, E., Rocha-Muniz, C. N., & Filippini, R. (2017). Understanding Auditory Processing Disorder Through the FFR. In N. Kraus, S. Anderson, T. White-Schwoch, R. Fay, & A. Popper (Eds.), The Frequency-Following Response: A Window into Human Communication (pp. 225–250). Springer International Publishing. 10.1007/978-3-319-47944-6_5

Sebastian-Gallés, N., Rodríguez-Fornells, A., de Diego-Balaguer, R., & Díaz, B. (2006). First-and Second-language Phonological Representations in the Mental Lexicon. Journal of Cognitive Neuroscience, 18(8), 1277–1291. 10.1162/jocn.2006.18.8.1277

Skoe, E., Burakiewicz, E., Figueiredo, M., & Hardin, M. (2017). Basic neural processing of sound in adults is influenced by bilingual experience. Neuroscience, 349, 278–290. 10.1016/j.neuroscience.2017.02.049

Skoe, E., & Kraus, N. (2010). Auditory Brain Stem Response to Complex Sounds: A Tutorial. Ear and Hearing, 31(3), 302–324. papers2://publication/uuid/07E666C5-A423-43C0-85EC-BA2067A7F59F

Song, J. H., Skoe, E., Wong, P. C. M., & Kraus, N. (2008). Plasticity in the adult human auditory brainstem following short-term linguistic training. Journal of Cognitive Neuroscience, 20(10), 1892–1902. 10.1162/jocn.2008.20131.Plasticity

The Jamovi Project. (2023). Jamovi 2.3. Retrieved from https://www.jamovi.org.

The Mathworks, I. (2019). MATLAB R2019b (p. Natick, Massachusetts).

Tomasello, M. (2000). Do young children have adult syntactic competence? Cognition, 74, 209–253.

Van Bezooijen, R. (1995). Sociocultural aspects of pitch differences between Japanese and Dutch women. Language and Speech, 38, 253–265.

Vouloumanos, A., & Werker, J. F. (2007). Listening to language at birth: evidence for a bias for speech in neonates. Developmental Science, 10, 159–164.

Weaver, I., Cervoni, N., Champagne, F., D’Alessio, A., Sharma, S., Seckl, J., Dymov, S., Szyf, M., & Meaney, M. (2004). Epigenetic programming by maternal behavior. Nature Neuroscience, 7, 847–854.

Werker, J. F., & Curtin, S. (2005). PRIMIR: A developmental framework of infant speech processing. Language Learning and Development, 1(2), 197–234. 10.1207/s15473341lld0102_4

Werker, J. F., & Hensch, T. (2015). Critical periods in speech perception: new directions. Annual Review of Psychology, 66, 173–196.

Werker, J. F., & Tees, R. C. (2005). Speech Perception as a Window for Understanding Plasticity and Commitment in Language Systems of the Brain. Developmental Psychobiology, 46(3), 233–251. 10.1002/dev.20060

White-Schwoch, T., Davies, E. C., Thompson, E. C., Woodruff Carr, K., Nicol, T., Bradlow, A. R., & Kraus, N. (2015). Auditory-neurophysiological responses to speech during early childhood: Effects of background noise. Hearing Research, 328, 34–47.

