## Supplementary Table 1 for "Exposure to bilingual or monolingual maternal speech during pregnancy affects the neurophysiological encoding of speech sounds in neonates differently"

,

^1^ Brainlab – Cognitive Neuroscience Research Group. Department of Clinical Psychology and Psychobiology, University of Barcelona (Catalonia, Spain)

^2^ Institute of Neurosciences, University of Barcelona (Catalonia, Spain)

^3^ Institut de Recerca Sant Joan de Déu, Santa Rosa 39-57, 08950 Esplugues de Llobregat (Catalonia, Spain)

^4^ BCNatal – Barcelona Center for Maternal Fetal and Neonatal Medicine (Hospital Sant Joan de Déu and Hospital Clínic), University of Barcelona (Catalonia, Spain)

^❖^ Both authors contributed equally to the preparation of the present work

*** Corresponding authors:**

*Carles Escera*

Brainlab - Cognitive Neuroscience Research Group

Department of Clinical Psychology and Psychobiology

University of Barcelona

P. Vall d'Hebron 171, 08035 Barcelona

Catalonia-Spain

*Jordi Costa-Faidella*

Brainlab - Cognitive Neuroscience Research Group

Department of Clinical Psychology and Psychobiology

University of Barcelona

P. Vall d'Hebron 171, 08035 Barcelona

Catalonia-Spain

**TABLES**

**Supplementary Table 1**

**Table 1. Descriptive statistics for MON** (n=53) **and BIL** (n=78) **groups in FFR parameters**: neural lag, Root-Mean-Square from pre-stimulus section, spectral amplitude at F_0_ and F_1_ peaks, Signal-to-noise ratio at F_0_ and F_1_ peaks.

| Measure | Mean | SD | Median | Q_1_ | Q_3_ | IQR | Minimum | Maximum |
| --- | --- | --- | --- | --- | --- | --- | --- | --- |
| **Neural lag** (ms) |  |  |  |  |  |  |  |  |
| Monolingual | 7.913 | 1.233 | 7.950 | 7.238 | 8.738 | 1.500 | 4.200 | 11.025 |
| Bilingual | 8.016 | 1.394 | 7.800 | 6.975 | 8.850 | 1.875 | 5.325 | 12.975 |
| **Pre-stimulus RMS** (nV) |  |  |  |  |  |  |  |  |
| Monolingual | 30.186 | 15.723 | 26.945 | 18.821 | 35.969 | 17.148 | 13.533 | 88.444 |
| Bilingual | 29.851 | 11.921 | 29.009 | 21.013 | 35.727 | 14.714 | 13.556 | 71.361 |
| **F_0_ Spectral Amplitude** (nV) |  |  |  |  |  |  |  |  |
| Monolingual | 10.015 | 5.184 | 9.042 | 6.495 | 12.270 | 5.775 | 1.503 | 26.110 |
| Bilingual | 9.186 | 4.924 | 7.709 | 6.063 | 11.236 | 5.173 | 2.568 | 23.960 |
| **SNR F_0_** |  |  |  |  |  |  |  |  |
| Monolingual | 4.607 | 3.512 | 5.245 | 2.589 | 7.377 | 4.788 | -4.351 | 10.770 |
| Bilingual | 3.200 | 3.741 | 3.967 | 1.294 | 5.504 | 4.210 | -6.739 | 10.224 |
| **F_1_ Spectral Amplitude /o/ section at 452 Hz** (nV) |  |  |  |  |  |  |  |  |
| Monolingual | 3.537 | 5.375 | 2.564 | 1.093 | 3.701 | 2.608 | 0.118 | 35.135 |
| Bilingual | 1.975 | 1.519 | 1.533 | 0.990 | 2.491 | 1.501 | 0.369 | 9.263 |
| **F_1_ Spectral Amplitude /a/ steady section at 452 Hz** (nV) |  |  |  |  |  |  |  |  |
| Monolingual | 2.118 | 1.373 | 1.863 | 1.354 | 2.619 | 1.265 | 0.279 | 7.869 |
| Bilingual | 1.866 | 1.314 | 1.731 | 0.946 | 2.434 | 1.488 | 0.242 | 9.405 |
| **F_1_ Spectral Amplitude /o/ section at 678 Hz** (nV) |  |  |  |  |  |  |  |  |
| Monolingual | 0.819 | 0.963 | 0.599 | 0.388 | 0.937 | 0.549 | 0.080 | 6.424 |
| Bilingual | 0.645 | 0.542 | 0.557 | 0.372 | 0.743 | 0.371 | 0.120 | 4.481 |
| **F_1_ Spectral Amplitude /a/ steady section at 678 Hz** (nV) |  |  |  |  |  |  |  |  |
| Monolingual | 2.519 | 5.948 | 1.063 | 0.555 | 1.918 | 1.363 | 0.167 | 38.348 |
| Bilingual | 0.929 | 0.726 | 0.759 | 0.603 | 1.353 | 0.750 | 0.061 | 5.068 |
| **SNR /o/ section at 452 Hz** |  |  |  |  |  |  |  |  |
| Monolingual | 0.516 | 5.435 | 1.928 | -1.678 | 4.406 | 6.084 | -18.893 | 6.388 |
| Bilingual | 0.599 | 3.643 | 1.262 | -1.514 | 3.772 | 5.286 | -9.345 | 6.043 |
| **SNR /a/ steady section at 452 Hz** |  |  |  |  |  |  |  |  |
| Monolingual | 0.913 | 3.448 | 1.231 | -0.718 | 3.214 | 3.932 | -8.834 | 6.078 |
| Bilingual | 0.161 | 4.116 | 1.360 | -2.556 | 3.284 | 5.840 | -13.965 | 6.196 |
| **SNR /o/ section at 678 Hz** |  |  |  |  |  |  |  |  |
| Monolingual | -1.112 | 4.395 | -0.174 | -4.008 | 2.002 | 6.010 | -14.753 | 6.236 |
| Bilingual | -1.008 | 3.538 | -0.748 | -2.852 | 1.766 | 4.618 | -10.866 | 4.321 |
| **SNR /a/ steady section at 678 Hz** |  |  |  |  |  |  |  |  |
| Monolingual | 1.431 | 4.305 | 2.744 | -0.975 | 4.597 | 5.572 | -14.170 | 6.660 |
| Bilingual | 0.142 | 4.792 | 1.802 | -2.724 | 3.661 | 6.385 | -16.269 | 5.982 |

SD = standard deviation. Q_1_ = first quartile (25th percentile). Q_3_ = third quartile (75th percentile). IQR = interquartile range.
